# SPEX: A modular end-to-end platform for high-plex tissue spatial omics analysis

**DOI:** 10.1101/2022.08.22.504841

**Authors:** Xiao Li, Ximo Pechuan-Jorge, Tyler Risom, Conrad Foo, Alexander Prilipko, Artem Zubkov, Caleb Chan, Patrick Chang, Frank Peale, James Ziai, Sandra Rost, Derrek Hibar, Lisa McGinnis, Evgeniy Tabatsky, Xin Ye, Hector Corrada Bravo, Zhen Shi, Malgorzata Nowicka, Jon Scherdin, James Cowan, Jennifer Giltnane, Darya Orlova, Rajiv Jesudason

## Abstract

Recent advancements in transcriptomics and proteomics have opened the possibility for spatially resolved molecular characterization of tissue architecture with the promise of enabling a deeper understanding of tissue biology in either homeostasis or disease. The wealth of data generated by these technologies has recently driven the development of a wide range of computational methods. These methods have the requirement of advanced coding fluency to be applied and integrated across the full spatial omics analysis process thus presenting a hurdle for widespread adoption by the biology research community. To address this, we introduce SPEX (Spatial Expression Explorer), a web-based analysis platform that employs modular analysis pipeline design, accessible through a user-friendly interface. SPEX’s infrastructure allows for streamlined access to open source image data management systems,analysis modules, and fully integrated data visualization solutions. Analysis modules include essential steps covering image processing, single-cell and spatial analysis. We demonstrate SPEX’s ability to facilitate the discovery of biological insights in spatially resolved omics datasets from healthy tissue to tumor samples.

## Introduction

Spatially resolved, highly multiplexed protein and RNA profiling of tissues at the cellular or sub-cellular level is essential for understanding the molecular components underlying tissue archi-tecture and function^1^. This fine characterization enables the measurement of cellular heterogeneity and cell-cell interactions in the spatial dimension which in turn brings us closer to identifying molecular determinants of tissue function. Hence, recently developed technologies, like IMC^2^, MIBI^3^, CODEX^4^ or MERFISH^5^, that allow for the acquisition of spatially resolved proteomic and transcriptomic states of individual cells and tissues, hold a great promise for advancing our understanding in many areas of cell and developmental biology. However, the widespread adoption of these technologies vitally depends on the development of analytical capabilities and infrastructure to readily extract biologically relevant conclusions from the complex data generated.

Spatially resolved omics data contain multiple layers of information which in turn are derived from imaging data and the molecular measurements associated with it. Examples of said layers constitute cell morphology, patterns of protein and transcript expression, cell neighborhoods or cell-cell communication at different spatial scales. To obtain this information, robust pre-processing and visualization methods are required as well as ways of analyzing the omics information and integrating it with the image derived properties^6^. The current software ecosystem developed in the past five years leverages several common analysis categories, including cell segmentation, clustering and spatial analytics, all required for meaningful interpretation of spatially resolved omics data^7,8^.

However, currently existing tools and custom analysis protocols provide only fragmentary solutions to the spatial omics analytical workflow^9,10^. None of them standing alone provide means for efficient data representation and pre-processing, interactive visualizations, and spatial relationship querying. As a result, most current end-to-end analytical strategies involve piecemeal workflows requiring data migration across numerous programs and coding proficiency. To our knowledge, the code-free platforms currently available are predominantly developed by commercial spatial omics platform vendors. These software solutions are user-friendly but often entail prohibitive licensing agreements and associated costs that hinder widespread use. Moreover, these commercial solutions demonstrate limitations in their applicability across various modalities or platforms.

Every imaging, transcriptomic, and proteomic data type has unique interpretation requirements, undercutting the effectiveness of platforms optimized for specific vendor data types. Therefore, the scientific community’s demand is twofold: for an open-source solution to democratize access and foster enhanced collaboration and reproducibility; and for a flexible, comprehensive tool that can be tailored to different modalities and platforms without compromising user accessibility and functionality.

With these current challenges in mind, we developed SPEX (Spatial Expression Explorer), a modular analytics platform that covers a broad span of essential methods required to analyze spatial omics data. SPEX is an application that could readily be deployed on cloud, or on a local workstation. It has a unique architecture that allows users to build data modality-specific customized pipelines from SPEX’s novel analytics developments or via plugging in external open source analysis modules. To support the needs of the cross-functional research community we designed SPEX in a way that one could operate it either from the user interface (code-free environment) or via more involved hands-on coding.

Here, as part of the SPEX software package, we implement extensible methods for image preprocessing, single-cell segmentation, segmentation post-processing, single-cell clustering, cell-cell co-occurrence, niche/neighborhood analysis, and spatially informed functional analysis in the form of differential expression analysis and pathway enrichment analysis. SPEX is integrated with the open-sourced and widely adopted image management system, OMERO^11^ and also leverages open-source spatial omics data visualization solution, Vitessce^12^, as a fully integrated element of the graphical user interface.

As a proof of concept, we show SPEX appropriately characterizes the spatial and molecular configuration of tonsil tissue. We further demonstrate the robustness of these methods in more heterogeneous tumor microenvironments represented in Pancreatic Ductal Adenocarcinoma (PDAC) and Triple Negative Breast Cancer (TNBC). Moreover, we extended the analysis to identify new patterns in tumor-immune microenvironment composition. Lastly, in a spatial transcriptomics dataset, we applied spatially-informed pathway analysis to a human lung cancer specimen and elucidated the pathways associated with immune attraction/avoidance in immune cells and tumor cells.

## Results

SPEX is a comprehensive spatial omics analysis platform implemented as a user-friendly web-based application with flexible plug-in analysis modules (Figure 1). SPEX provides both infrastructure and quantitative analysis methods that allow for efficient storage (Figure 1a), manipulation and interactive visualization of spatial expression data (Figure 1e). SPEX has an easy to use graphical user interface which allows the wider research community to build analytical pipelines and visualize high dimensional spatial data.

**Figure 1.**
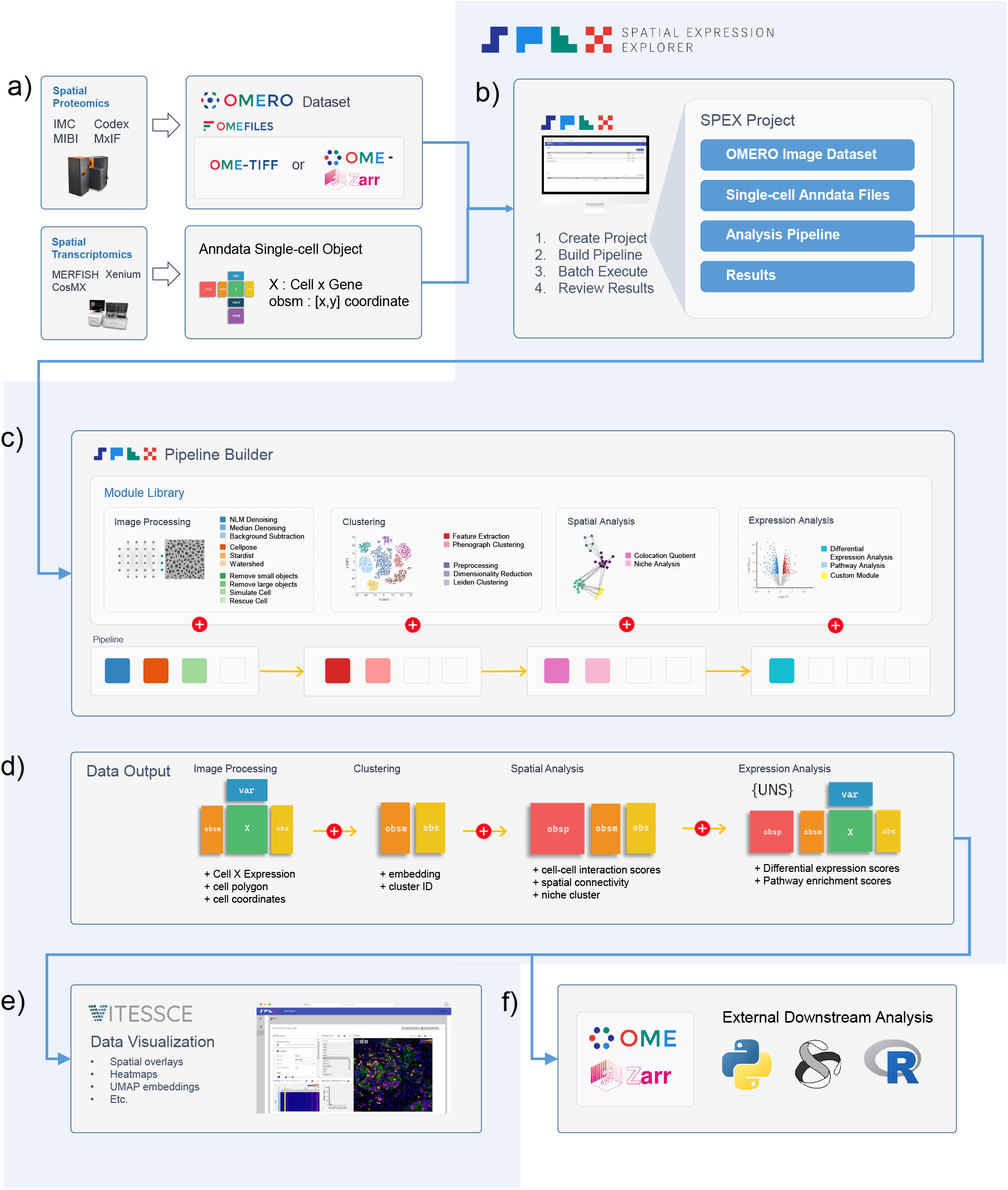
Graphical depiction of the Spatial Expression Explorer (SPEX) platform and analytical workflow. **(a)** Data can be input into SPEX as images or single-cell objects. Image data is managed and served into SPEX via the OMERO image management system. **(b)** Analytical projects in SPEX are executed as a 4 step process - create project, build pipeline, batch execute and review results. Projects are containers for input data, editable pipelines and study output data. **(c)** A modular pipeline builder is used to assemble an analytical routine. Modules are selected from the module library, covering image processing, segmentation, clustering, spatial analysis and expression analysis. The individual modules are shown as multi-colored rounded squares and can be added to the pipeline in a graphical manner. **(d)** Analysis data generated by the pipeline are stored as an Anndata object. As the pipeline proceeds, new elements are added to the Anndata object. **(e)** Processed datasets packaged as Anndata ZARR files can be reviewed using an integrated Vitessce dashboard. **(f)** Output data is compiled as an Anndata ZARR can be exported for downstream analysis in a variety of platforms.

### SPEX Analytical Modules

For spatial proteomics modalities, SPEX can start with raw imaging data or single-cell objects with spatial coordinates. For spatial transcriptomics modalities, SPEX requires a single-cell object with spatial coordinates as inputAnalysis can then be performed on these assets by leveraging a wide range of algorithms that can be categorized into four primary domains-image processing (spatial proteomics), single-cell clustering, cell-cell interaction spatial analysis and spatially informed differential expression analysis and pathway analysis.

#### Image Processing Modules

SPEX includes a modular pipeline to facilitate tissue-based single-cell segmentation (Figure 1c). This generalized pipeline aims to accommodate a wide range of high-dimensional imaging modalities such as IMC, MIBI, cyclic IF and spatial transcriptomics . Image processing is executed in a 4 step sequence with the ultimate goal of generating a cell by expression matrix in Anndata format for downstream single-cell and spatial analysis. These steps include image preprocessing, single-cell segmentation, post-processing and feature extraction. Each step contains a selection of modules which can be linked together to address the particularities of a given image set. The adaptability of the pipeline was demonstrated by analyzing imaging data coming from a variety of high dimensional imaging modalities. Each imageset presented unique characteristics; varying in resolution, signal to noise and structural attributes.

#### Clustering Module

To enable the analysis of the high-plex omics data associated with the images, SPEX provides single-cell clustering modules that cover both intensity-based proteomics and count-based transcriptomics single-cell inputs. For transcriptomics, we rely on the suite of proven methods offered by the Pegasus package^13^. For proteomics, we also provide the widely adopted graph-based clustering algorithm, Phenograph^14^.

#### Spatial Analysis Module

Tissue architecture can be complex, with cell types forming spatial patterns that define particular domains where they might exert very different functions owing to the distinct local cellular context. These functions might be reflected in the associated gene or protein expression patterns. SPEX implements the Colocation Quotient (CLQ) module, as detailed in the methods section, which integrates spatial coordinates of the identified cells with their respective cell types to stratify them into co-location or avoidance pattern between cell types. Further, given a set of spatially co-occurring cells, it is natural to ask whether these co-occurrence patterns are repeated throughout the tissue, and what comprises such patterns or spatial niches. Clustering spatial co-occurrence features (in an unsupervised fashion) facilitates the identification of such spatial patterns or niches.

#### Spatial expression analysis Module

The functional state of cells and/or their molecular action may be influenced by their spatial organization in tissue. This can be due to inclusion in higher order functional structures or cell-cell mediated interactions. To facilitate quantification of spatially informed expression, SPEX includes both differential expression analysis and pathway analysis. These modules can take spatially informed cell categories as input

### SPEX User Interface

The SPEX graphical user interface is designed to minimize spurious user options and visual overhead in an effort to increase platform intuitiveness. At a high-level the SPEX interface can be divided into 3 primary sections organized as sequential steps-project creation,data loading, and analysis development (Figure 2). The analysis section contains sequential sub-sections covering pipeline building, batch execution and data visualization (Figure 2c).

**Figure 2.**
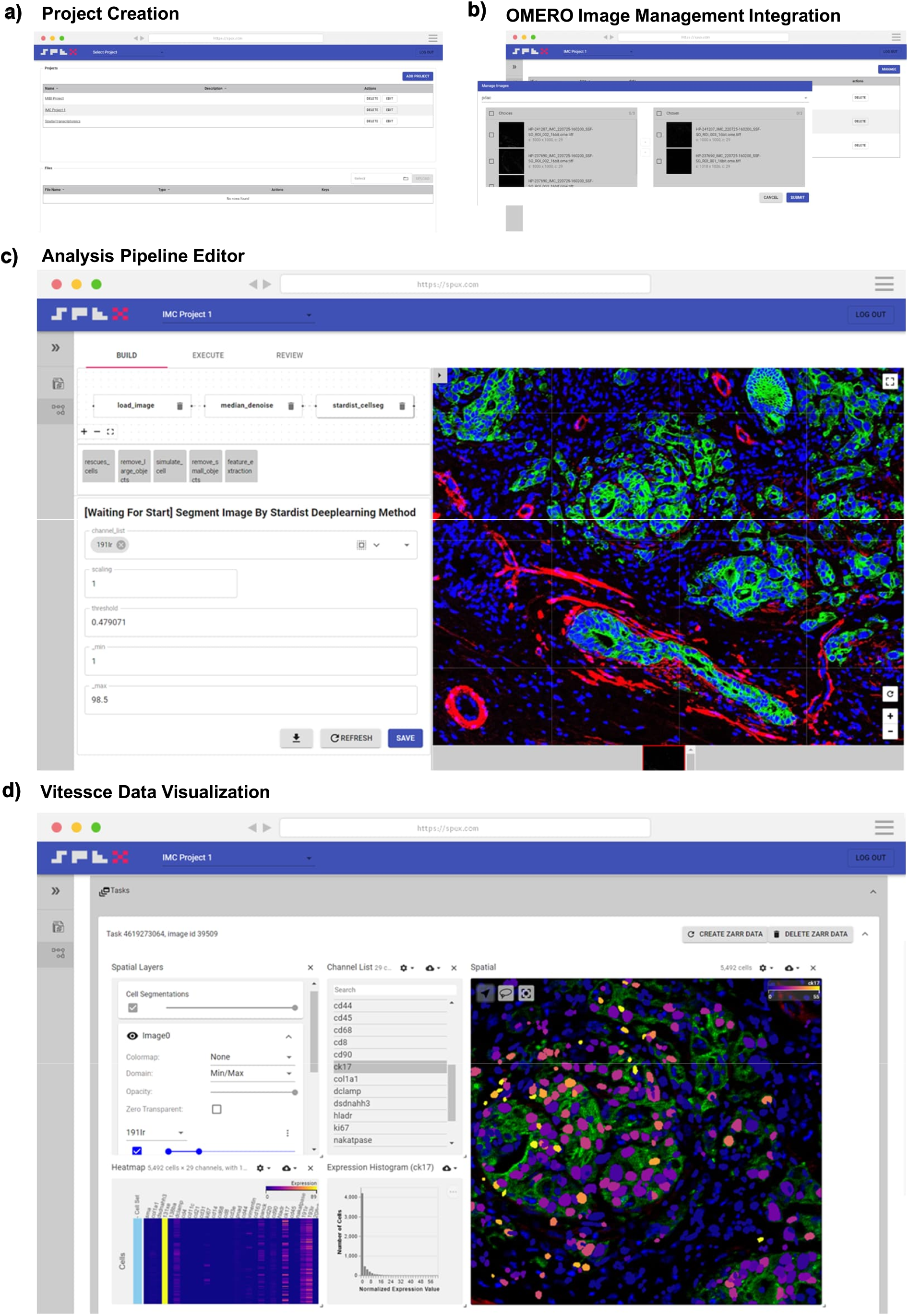
SPEX Graphical User Interface. **(a)** SPEX project creation user interface. Projects are containers for study data. **(b)** SPEX-OMERO integration highlighting navigation and selection OMERO hosted datasets and images within SPEX. **(c)** SPEX pipeline builder user interface. An intuitive 3 step process of building, executing and reviewing the analysis displayed at the top of the interface. The user moves through each step in sequence. In the build section the user is able to graphically build an analysis pipeline by selecting and chaining modules in sequence. A unique parameter section is displayed for each module. **(d)** Fully integrated Vitessce dashboard for visualizing results in spatial context.

After logging into SPEX, the first screen encountered by users is the SPEX project creation interface (Figure 2a), where projects, serving as containers for study data, are initiated. The user can define a name and description for the project for future retrieval. This project can then be accessed and populated with data to be analyzed. SPEX supports one of three data formats - OMETIFF, OMEZARR or H5AD (Anndata). OMETIFF and OMEZARR represent supported multiplex image formats while H5AD represents the supported single-cell data format. The integration with OMERO (Figure 2b) allows seamless navigation and selection of OMERO-hosted image datasets within SPEX. Once a project is populated with image or single-cell data, the user can define an analysis pipeline. Several pipelines can exist for any given project if the user is interested in prototyping different analysis workflows. To develop an analytical workflow, users engage with the SPEX pipeline builder interface (Figure 2c). The user progresses In the build section enabling the graphical construction of analysis pipelines. In this phase, modules are selected from a library and dragged into a visual pipeline map (Figure 1c). The selection of modules are dependency-informed, where new modules can only be added to modules that generate pre-requisite data, thus eliminating the potential for erroneous pipeline construction. In the execution page of the pipeline builder, the user can send the data for batch processing. The status of the analysis jobs will be displayed for all images in the queue. When complete, an aggregated Anndata single-cell object housing data from all images will be generated and available for local download as a ZARR file. This ZARR file can be loaded into other external analysis routines in Python or R (Figure 1f). The final step in the pipeline building interface is data visualization. Here, users can visualize analytical readouts using a Vitessce spatial omics dashboard displaying common plots such as scatterplots of UMAP embeddings, cluster by expression heatmaps, histograms, spatial heatmaps and more. All visualization panes are interactive and have cross-filtering functionality (Figure 2d). This cohesive workflow ensures a user-friendly and efficient analysis experience within the SPEX environment.

### SPEX workflow on spatial proteomics data

#### SPEX identified structural composition of Tonsil

In the previous sections, we illustrated the capabilities of the different modules that integrate SPEX. To demonstrate that SPEX analysis modules generate biologically meaningful results, we validated the SPEX workflow by analyzing a sample from human tonsil tissue, which has a well-defined cellular organization^15–17^. To achieve this, a 4µm-thick section of formalin-fixed, paraffin-embedded (FFPE) tonsil tissue was stained with a panel of Imaging Mass Cytometry (IMC) antibodies listed in Supplementary Table 1. The antibody panel was designed to simultaneously characterize the composition of the immune compartment, the spatial relationship between immune cells and stromal cells, and the interactions among cell subsets.

The stained tonsil sample was imaged with the Hyperion Imaging System and then pre-processed and analyzed using a SPEX pipeline utilizing most of SPEX modules (Figure 3a). This pipeline applied to the tonsil tissue IMC images started with per cell segmentation using StarDist based on the nuclear histone H3 channel. Then, Phenograph was used for clustering the segmented cells according to their median marker expression levels of the different IMC channels. Clustering outcomes were visualized using UMAP, and cluster labels were assigned based on the IMC marker measurements as well as positional data (x and y coordinates) of cells revealing 3 well-defined lobes representing epithelial, follicular and paracortical tissue compartments distinguished primarily by expression of cytokeratin, Ki67, and CD8, respectively (Figure 3b), consistent with the composition of cells in these compartments. Within the individual UMAP regions, the location of closely related cell populations correlates with biologically relevant parameters.For example, we note distinct populations representing well characterized functional zones of tonsil germinal centers. Germinal center light and dark zones can be distinguished across an axis of Ki67, BCL2 and CD45 expression.

**Figure 3.**
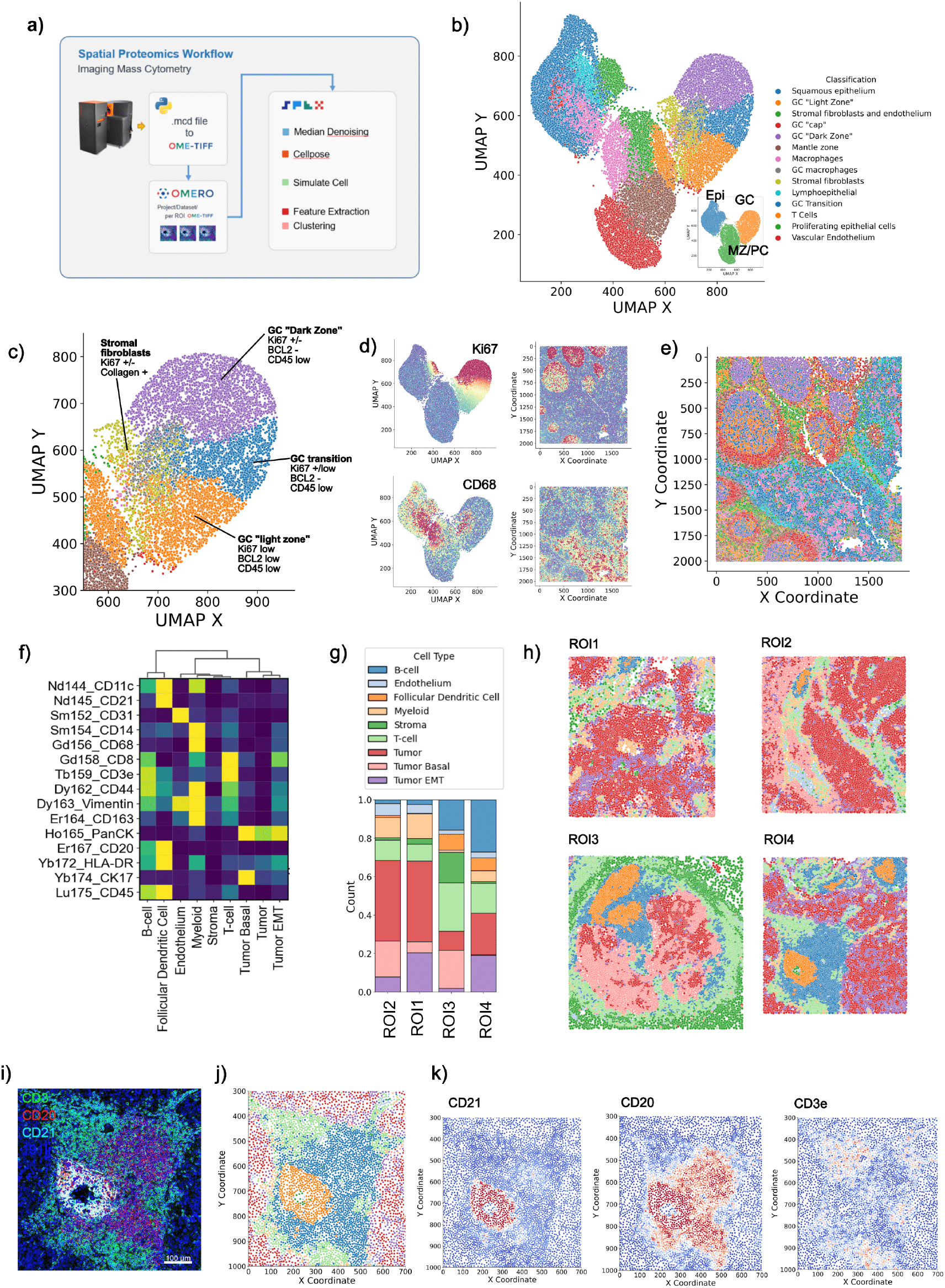
SPEX robustly identified structural composition of Tonsil and PDAC tissue in Imaging Mass Cytometry modality. **(a)** Graphical dataset description and SPEX spatial proteomics workflow modules highlighted in this figure **(b)** UMAP embedding of Tonsil single-cell data colored by pathologist annotated cell type. Inset shows primary high order Tonsil structures (EPI=Epithelial, GC = Germinal Center, MZ/PC= Marginal Zone/Paracortex). **(c)** Detail of the Germinal Center region of UMAP with pathologist annotated functional zones. **(d)** Protein expression of Ki67 and CD68 color mapped to UMAP embedding and spatial domain. Warm colors denote high expressing regions **(e)** The histologic tonsil image is color-coded by 14 structural hierarchy categories identified by the marker expression pattern. **(f)** PDAC single-cell clustering expression heatmap **(g)** Cell type composition of 4 individual IMC ROIs **(h)** Spatial map of clustered cell types across 4 PDAC ROIs **(i)** Multiplex IMC image showing detail of Tertiary Lymphoid Structure (TLS). CD3= Green, CD20 = Red, CD21=Cyan **(j)** Detail spatial map of cell types in a TLS region **(k)** Spatial distribution expression heatmap of CD21, CD20, and CD3e.

We applied this same SPEX workflow of single-cell segmentation, feature extraction, data normalization, and single cell phenotyping to four regions of interest in a pancreatic adenocarcinoma (PDAC) patient tumor, revealing major cell types by Phenograph, including stromal cells, T cells, myeloid cells, B cells, FDC cells, and a myriad of tumor cell states (Figure 3f). Compositional analysis of these cell states per ROI revealed two of the four ROIs encompassed robust B cell and FDC cell frequencies (Figure 3g) in a spatially co-localized pattern (Figure 3h). Upon further spatial interrogation of the molecular expression of CD3, CD20, and CD21 in these cells by image overlays (Figure 3i) and spatial heatmaps (Figure 3j,k, l) they appear to compile a Tertiary Lymphoid Structure (TLS).

#### SPEX Spatial Proteomics Pipeline Enables Novel Single Cell Spatial Insights

Next we analyzed a public Multiplex Ion Beam Imaging (MIBI) dataset^3^ to demonstrate the single cell spatial analysis methods of the SPEX pipeline (Figure 4a). This imageset consisted of 41 field of view images with 36 channels. Being a non-optical mass detection platform, it is common for raw images to include substantial noise when compared to traditional optical acquisition platforms. Therefore, median denoising was executed on channels utilized for segmentation (dsDNA, H3K9ac, H3K27me3), channels were then merged and segmented for single-cells using the Cellpose deep learning model^18^. Following cell phenotyping by unsupervised clustering these SPEX modules generated cell calls that correctly match their molecular profiles and the published calls by the original work^3^ (Figure 4b-d). Through arranging the tumors from low to high immune cell frequency of total cells, we observe an increased diversity of immune cell types within the sample where samples with a lower density of immune cells are dominated by macrophages and samples with higher density of immune cells have a mixed composition of lymphocytes, myeloid cells and antigen presenting cells (Figure 4d). To further interrogate the spatial relationship between this variable immune compartment and the tumor cells, we employ the Colocation Quotient (CLQ) method under the SPEX Spatial co-occurrence module. Briefly, the CLQ measures the co-occurrence or avoidance of cell type pairs by looking at the local density of a target cell type at a fixed radius from each cell of the sample belonging to a reference cell type. That way, one can, for instance, query if target immune cells are co-locating or avoiding tumor cells. We applied the CLQ module with the immune cell::tumor cell pair here in samples that had more than 10% immune cells (non-desert samples) revealing two major groups of samples, those with high immune::tumor cell spatial enrichment, and those with low immune:tumor cell enrichment (Figure 4e). These identified inflamed and excluded groupings aligned well with the visual representation of immunophenotyping presented in the images (Figure 4f).

**Figure 4.**
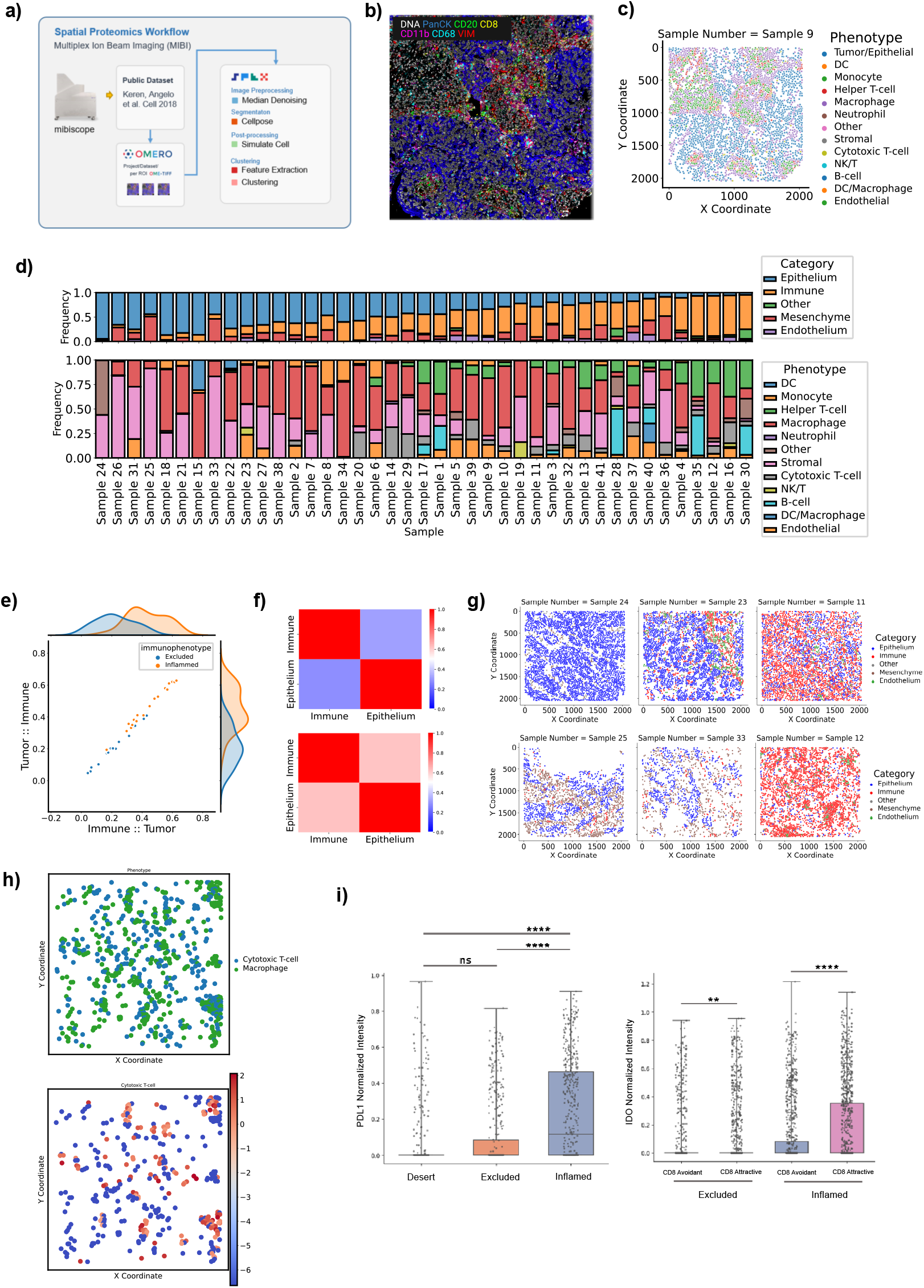
SPEX spatial analysis modules reveal immune cell state relationships to tumor:immune spatial architecture in a TNBC MIBI dataset. **(a)** Graphical dataset description and SPEX spatial proteomics workflow modules highlighted in this figure. **(b)** Representative image of TNBC patient tumor from the Keren cohort showing color overlays of DNA (white), PanCytokeratin (PanCK, blue), CD20 (green), CD8 (yellow), CD11b (pink), CD68 (cyan) and Vimentin (VIM, red), scale bar = 100 um. **(c)** Cell phenotype map corresponding to B showing 13 major cellular phenotypes by colored points. **(d)** Frequencies of major cell lineages (of all cells, top) and immune cell types (of total immune cells, bottom) are shown as stacked bar plots for each patient. **(e)** Scatterplot showing results of SPEX CLQ analysis module camparing the immune::tumor spatial enrichment vs tumor:immune spatial enrichment in each patient that had more than 10% immune cells (not desert). Based on these data patients are assigned to an immune-inflamed (orange) class or an immune-excluded (blue) class using a GMM model. **(f)** Heatmaps showing the tumor:immune, tumor:tumor, immune:tumor, and immune:immune CLQ results in immune-excluded patients versus immune-inflamed patients. **(g)** Representative cell phenotype maps of two immune-desert tumors, two immune-excluded tumors, and two immune-inflamed tumors, showing the location of tumor (blue), immune (red), mesenchymal (brown), endothelial (green), and other (grey) cells. **(h)** A cell phenotype map (top) showing cytotoxic T cells (blue) and macrophages (green) is shown above a heatmap of the macrophage cells, colored by their CLQ Macrophage:Tcell spatial enrichment score. **(i)** Boxplot showing the normalized expression of PDL1 in macrophages in immune-desert, immune-excluded, and immune-inflamed tumors, asterisks denote significance in kruskal-wallis test, **** P < 0.001, ns = not significant. Boxplot showing the normalized expression of IDO1 in macrophages that are either CD8-Tcell-avoidant (CLQ < 0.5) or CD8-Tcell-attractive (CLQ> 1.5) in immune-excluded vs immune-inflamed tumors, Mann-whitney test, asterisks denote significance: ** *P* < 0.01, **** *P* < 0.001.

Having defined the tumor immune phenotypes with CLQ analysis as desert, excluded, or inflamed tumors, we explored potential gene expression patterns yielded by this classification. Of clinical relevance, we observed a significant increase in PDL1 expression by tumor-associated macrophages in inflamed tumors suggesting the tumor-immune spatial relationships may represent informative biomarkers of immunotherapy potential (Figure 4h)^19,20^. We then further leveraged the CLQ module to interrogate macrophage phenotypes in these tumors by computing their pairwise spatial enrichment with all other cell types. Interestingly, this combination of CLQ and expression analysis revealed that macrophages that are spatially enriched (attractive) with CD8+ Cytotoxic T Cells (Figure 4g) had significantly higher IDO1 expression compared to macrophages that are spatially isolated (avoidant) from these T Cells (Figure 4i). This relationship was observed in excluded tumors and further enriched in inflamed tumors, suggesting that in addition to an enhanced potential for PDL1 antagonists in this inflamed subset of TNBC patients, IDO1 may be an immunosuppressive target of interest to alleviate local macrophage suppression of T cell activity^21^.

### SPEX workflow on spatial transcriptomics data

The availability of spatial resolution enhances the meaningful interpretation of cellular interactions and signaling events. The CLQ module calculates descriptive statistics of spatial avoidance and attraction at both the single-cell and cell cluster level. Gene and pathway expression states can be correlated with these statistics to provide insight into the effects of spatial co-occurrence on the transcriptome.

As an example, we used the CLQ method to stratify cells in a human lung cancer MERFISH sample by attraction or avoidance to each cell type (Figure 5c). We investigated how proximity to T cells modulates inflammatory and cell proliferation pathways (Figure 5d). We noticed up-regulation of NFKb pathway genes and down-regulation of Trail and TNFa genes in epithelial cells, up-regulation of EGFR genes and down-regulation of TNFa genes in myeloid cells, and down-regulation of p54 genes in endothelial cells, when split by proximity to T cells. Furthermore, the integration of the CLQ with omics data allowed us to identify genes that are differentially expressed in the proximal presence or absence of another cell type. As an example, we assessed how gene expression differs between T cells that are classified as “attractive” to or “avoidant” of Epithelial cells (as determined by CLQ analysis).

**Figure 5.**
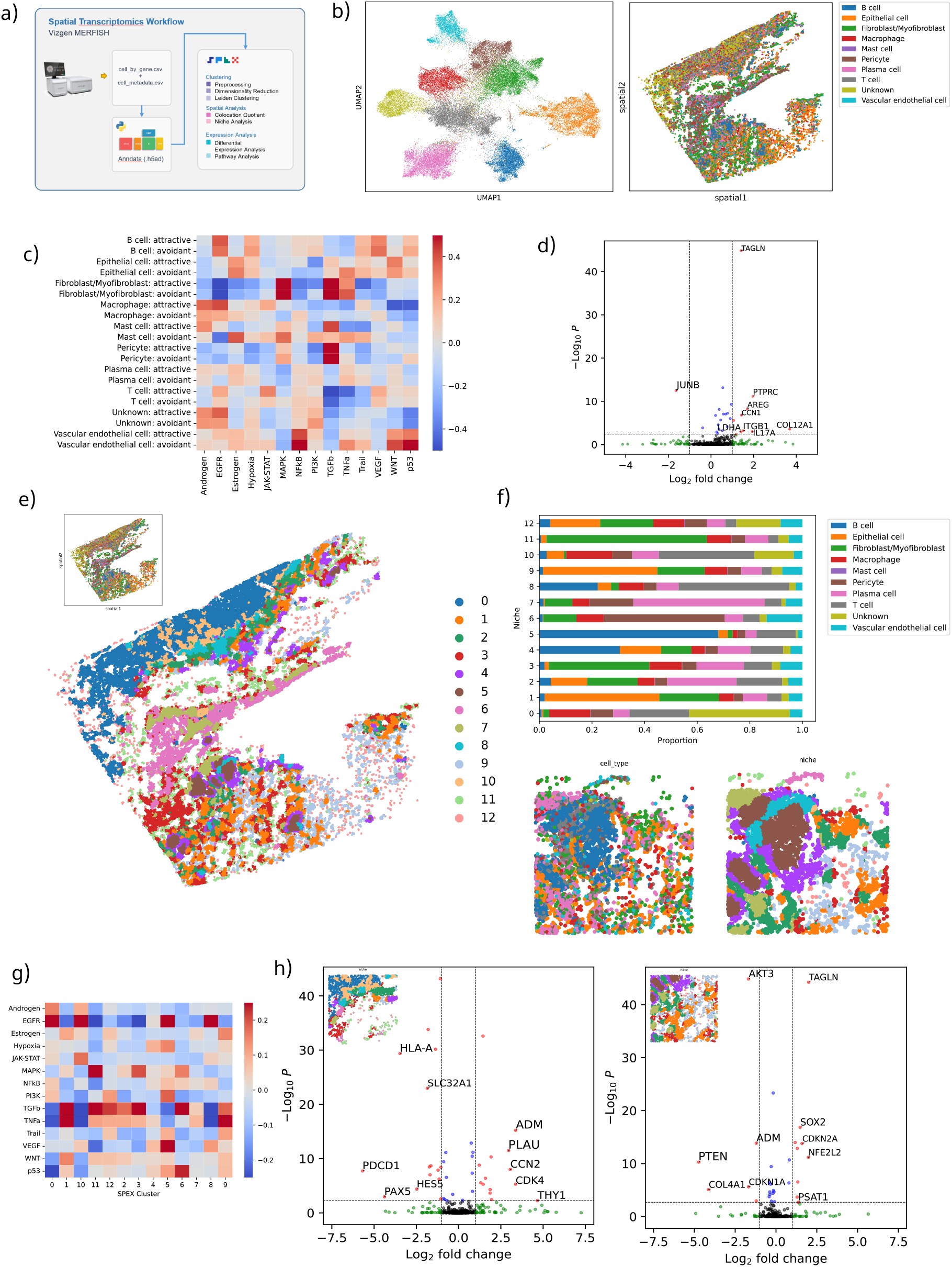
SPEX spatial transcriptomics analysis identifies immune cell niches. **(a)** Graphical dataset description and SPEX spatial transcriptomics workflow modules highlighted in this figure. **(b)** Cell phenotype map in both gene expression (UMAP, left) and spatial (right) coordinates. **(c)** PROGENy pathway enrichment scores for each cell type cluster, split by colocalization with T cells. Colocalization was determined using the SPEX CLQ analysis module. Scores were calculated by fitting a multilinear model to the gene expression of each cell given the pathway weights per gene. The average score per cluster is shown here. **(d)** Volcano plot showing differentially expressed genes for T cells based on colocalization with epithelial cells. **(e)** Spatial map of cell niches from (a). **(f)** Graphical dataset description and SPEX spatial transcriptomics workflow modules highlighted in this figure. **(g)** Graphical dataset description and SPEX spatial transcriptomics workflow modules highlighted in this figure. **(h)** PROGENy pathway enrichment scores for each spatial niche. **(i)** Volcano plot showing differential expression between T cells in spatial niche 10 vs spatial niche 8 (B-cell enriched) (left) and volcano plot showing differential expression between epithelial cells in spatial niche 9 (macrophage enriched) vs spatial niche 1 (right).

The CLQ method is limited to pairwise relations and does not take into account higher order interactions with additional cell types within potential functional communities composed of multiple cell types. We acknowledge that spatial patterns exist beyond pairwise spatial correlations and as such we introduce the next analysis module, SPEX niche analysis. This module identifies cell communities by determining the neighborhood composition for each cell in the image at a given radius and then clustering all the different neighborhoods to identify cell communities.

In the lung tumor MERFISH sample, the resulting analysis demonstrated the presence of cell type communities that define regions of interest in the image with potentially different biology. This allowed us to extract further information than the mere projection of the cell types identified by transcriptomics back into the image (Figure 5b). SPEX communities, when highlighted in the tissue slide, allow for the identification of potential structures (Figure 5e). For example, SPEX communities bring out aggregates of B cells surrounded by layer comprised of a combination of B cells, epithelial cells and T cells potentially defining TLS (Figure 5f). Beyond the spatial information, the clusters themselves can be characterized using pathway analysis. Pathways will be present as a function of the cell types involved in the community and their contextual transcriptomic state, In the present case, TGF has larger activity in communities containing more fibroblasts (Figure 5g).

The SPEX community analysis module can also be leveraged to look at specific cell types in the context of a particular microenvironment. More concretely, differential gene expression analyses for the same cell type across clusters can further identify context dependent expression patterns (Figure 5h). We highlight how the T cells tend to express more CD8A in SPEX community 8 (B-cell associated) compared to 10 potentially indicating a differentiation towards more cytotoxic T cells in that com-munity. Tumor cells also show some interesting patterns, when associated with stroma they express more HLA and CYR61, involved in angiogenesis. Overall, our community module generates hypotheses about regions of the tissue where potential contextual phenotypes are observed and guides the further exploration of the data.

## Discussion

Cells in complex multicellular organisms are hierarchically arranged into tissues that are in turn organized into organs, and so on. At each level of organization, structure is closely related to function. In tissues, both during and after development, structure is governed by different gene expression programs that in turn depend on the resulting architecture of the cellular microenvironment experienced by the individual cells composing the tissue. This suggests that if we can measure gene and protein expression of individual cells in a given tissue and efficiently incorporate spatial cell arrangement information into these measurements, we would be able to discern the molecular basis of tissue function and dynamics.

Currently, the fast pace of the development of spatial transcriptomics and proteomics^22^ is bringing us closer to fulfill this scientific program. Nevertheless, the data generated by these methodologies is very rich in information and its successful analysis and interpretation depends on the existence of computational pipelines. Most of the solutions provided by the scientific community are libraries for use by computational biologists which expose the need for a graphical user interface application to enable the direct interaction with the data from other domain experts such as pathologists or biomedical scientists. Although there are commercially licensed platforms that offer more accessible graphical interfaces, their application is restricted due to high costs, license limitations, and lack of generalizability across different modalities or platforms. This highlights the dual need for an open-source solution and a modular, easily adaptable tool.

In response to this requirement, we presented in this work, SPEX, a powerful toolkit that offers a graphical user interface for analysis of spatially resolved omics data. The unique modular design of SPEX provides flexibility, enabling users to create bespoke data analysis pipelines. Users can directly interact with data through the graphical interface, significantly easing the process of data analysis and interpretation by eliminating the need for low-level coding.

The SPEX platform demonstrates flexibility in analyzing diverse biological data types, illustrating both its breadth of functionality and its modality-agnostic adaptability. For IMC data, we leveraged methods in SPEX to establish an end-to-end workflow to systematically characterize the spatial organization and molecular features of the highly structured microenvironment of human tonsil tissue. Additional application extended to analyzing regions of interest in a human PDAC dataset wherein we characterized TLS positioned within a heterogeneous tumor microenvironment.

Our exploration continued in other high dimensional imaging modalities where we leveraged the CLQ method to quantify local cell-cell interaction in a TNBC MIBI dataset. The outcomes were not only consistent with previously reported results by orthogonal methods but also provided novel insights into the relationship between spatial proximity and function of macrophage populations present in varying microenvironment tumor-immune phenotypes.

Finally, we explored the ability of SPEX methods to tackle deeper data analysis tasks by analyzing a NSCLC spatial transcriptomics dataset. We integrated gene expression analysis with spatial information by leveraging the SPEX niche analysis module. We identified microenvironment domains within the tumor tissue where particular transcriptional programs are more active. Across all these use cases, SPEX showed its potential to discover and characterize informative biological programs from spatially resolved omics data.

We are aware that there are still many open questions that warrant the development of analysis modules. Our analyses indicate that “ambient RNA” diffusion in the slide is a problem of MERFISH samples that goes beyond the need to improve the cell segmentation step. This is an example of many possibilities for future development and, therefore, we hope that the scientific community will embrace this software tool and expand upon its modules.

While we foresee immediate utility of the current SPEX version in the emerging field of spatially resolved omics, we recognize the need to enhance its capabilities for the imaging processing of whole-slide imaging data and spatial transcriptomics modalities. In the forthcoming years, the field of spatially resolved omics is expected to progress towards generating multi-modal readouts, encompassing transcriptomics, proteomics, TCR and BCR repertoire, epigenomics, and metabolomics, all from the same tissue slide^22,23^. Therefore, we present SPEX which is available at https://github.com/Genentech/spex, not merely as a static tool, but as an adaptable, and extensible open-source solution, directly catering to the diverse and rapidly evolving needs of the spatial omics research community.

## Methods

### Data Overview

To demonstrate SPEX capabilities, we leverage both newly generated and published datasets. The published datasets include raw MIBI data for a cohort of 41 TNBC patients that was downloaded from https://www.angelolab.com/mibi-data^3^.

Control Tonsil Tissue Antibody staining for imaging mass cytometry was performed on 4µm-thick, formalin-fixed, paraffin-embedded tissue sections mounted on SuperFrost PlusTM glass slides. Antibodies to cytokeratin (cat. no. ab80826), CD103 (cat. no. ab221210) and PD-L1 (cat. no. ab226766) were purchased as purified, carrier-free formulations from AbCam and, using the Maxpar X8 Antibody Labeling Kit (Fluidigm Inc., South San Francisco, CA), conjugated to 176-Yb (cat. no. 201167A), 153-Eu (cat. no. 201153A) and 150-Nd (cat. no. 201150A), respectively, per manufacturer’s protocol. Remaining antibodies were purchased as conjugated from Fluidigm. To stain, tissue was first deparaffinized and rehydrated. Next, pH 9.0 antigen retrieval was used followed by incubation in Superblock at room temperature for 30 minutes. Samples were incubated with a staining panel comprised of 16 antibodies (see Supplementary table1) overnight at 4C in a humidity chamber. Samples were washed four times in 0.1% TritonX/PBS for four minutes each followed by two 4 minute washes in PBS at room temperature. Samples were then incubated with Ir-Intercalator dye for 30 min at room temperature, washed in water for 5 min and air-dried. Regions of interest (ROIs) measuring 1800 x 2000 um each were selected from the slide and acquired at 200 Hz. Data were exported as MCD files and visualized.

For PDAC Imaging Mass Cytometry samples, slides were first baked at 70°C for a minimum of 30 minutes to remove all visible wax. They were then subjected to a deparaffinization and rehydration series using xylenes, alcohol, and water in an autostainer. Antigen retrieval was subsequently achieved using the EZ-Retrieve^®^ System, and the slides were cooled to about 74°C in EZ-AR1 solution. After a 10-minute wash in Maxpar Water, the samples on the slide were encircled with a PAP pen and then blocked with a freshly prepared 3% BSA in Maxpar PBS solution. For antibody staining, an overnight MCA cocktail specific for the assay was prepared. The slides were incubated overnight with this mixture at 4 °C and given a secondary antibody for 30 minutes following a rinse in Maxpar PBS. Counterstaining involved washing slides with 0.2% Triton X-100 in Maxpar PBS followed by rinses in Maxpar PBS. Slides were then incubated in a diluted Intercalator-Ir solution. Finally, slides were washed in Maxpar Water, and air-dried overnight at room temperature.

MERFISH Lung Tumor Sample Samples from patients with human lung cancer were snap frozen and preserved in optimal cutting temperature (OCT) compound and cut into 10 µm thick on a cryostat at -20°C and placed on MERSCOPE Slide (Vizgen 20400001). The tissue slices were fixed with 4% paraformaldehyde in 1x PBS for 15 minutes, washed three times with 5 mL 1XPBS and incubated with 70% ethanol at 4 C overnight for tissue permeabilization. Samples were then stained for cell boundary using Vizgen’s Cell Boundary Kit (10400009), and later hy-bridized with a custom designed MERSCOPE Gene Panel Mix consisting of 484 genes that assess the different cell types (tumor, stroma, immune cells, etc) and key cell signaling and activity marker (Vizgen 20300008) at 37°C incubator for 36-48 hours. Following incubation, the tissues were washed with 5 mL Formamide Wash Buffer at 47°C for 30 minutes, twice and embedded into a hydrogel using the Gel Embedding Premix (Vizgen 20300004), ammonium persulfate (Sigma, 09913-100G) and TEMED (N,N,N’,N’-tetramethylethylenediamine) (Sigma, T7024-25ML) from the MERSCOPE Sample Prep Kit (10400012). After the gel mix solution solidified, the sam-ples were cleared with Clearing Solution consisting of 50uL of Protease K (NEB, P8107S) and 5mL of Clearing Premix (Vizgen 20300003) at 37°C overnight. After removing clearing solution, the sample was stained with DAPI and Poly T Reagent (Vizgen 20300021) for 15 minutes at room temperature, washed for 10 minutes with 5ml of Formamide Wash Buffer, and then im-aged on the MERSCOPE system (Vizgen 10000001). A fully detailed, step-by-step instruction on the MERFISH sample prep the full protocol is available at https://vizgen.com/resources/fresh-and-fixed-frozen-tissue-sample-preparation/. Full Instrumentation protocol is available at https://vizgen.com/resources/merscope-instrument/.

### SPEX modules

The SPEX graphical user interface provides essential components to facilitate comprehensive analysis of spatial omics images and datasets SPEX supports end to end modules for spatial proteomics - covering both image processing and downstream single-cell spatial analysis. Since the vast majority of single-cell spatial transcriptomics platforms output cell by transcript files with spatial coordinates, SPEX will also directly ingest a cell by transcript Anndata file (.h5ad) for downstream clustering and/or single-cell spatial analysis.

#### Image processing

Image processing is executed in a 5 step sequence with the ultimate goal of generating a cell by expression matrix. These steps include-image loading, image preprocessing, single-cell segmenta-tion, post-processing and feature extraction. Each step contains a selection of modules which can be linked together to address the particularities of a given image set.

#### Image loading

The image loading step contains a single module which supports OME-TIFF or OME-ZARR. This multi-dimensional open format provides a well-structured metadata header that can accommodate a wide range of image and acquisition system information. A number of open source conversion pipelines are available in the event proprietary microscopy formats need to be converted to OMETIFF or OME-ZARR. Within SPEX, the AICSimageio python library (https://github.com/AllenCellModeling/aicsimageio) is used read these open formats in as in-memory multi-channel Numpy arrays SPEX also supports the loading of h5ad files, which are a data format for AnnData objects.

This allows SPEX users to utilize h5ad files as data sources, fter loading, this data can be processed and analyzed in the SPEX pipeline, for those modules that provide this capability.

The preprocessing step includes optional modules to denoise images and/or enhance pixel information to facilitate single-cell segmentation. These modules include-global background correction, median filter denoising, and Non-local means (NLM) denoising. All modules were developed using the Scikit-image python library. To accommodate the potentially large channel dimension of images, channels are processed in parallel with use of the apply *parallel()* function in SCIKIT image, a wrapper for DASK map *blocks()* function. This routine allows application of the denoising functions on chunked arrays which are efficiently distributed across computing cores.

In the global background correction module, background signal, as captured in one channel, can be subtracted from all other channels. For example, signal in an autofluorescent or a detector noise channel can be subtracted from channels that house specific molecular markers. Within the defined background channel, OTSU thresholding^24^ is applied to create a positive pixel binary mask. This pixel mask is then intensity-scaled by a user-defined correction factor. This correction mask is then subtracted from the other channels.

Median denoising is a common image processing technique which can preserve edge information while suppressing unstructured noise. For any channel, the filter replaces a pixel’s value with the median of its local neighborhood. The size of the local neighborhood is defined by the filter kernel size, an argument of the function.

Non-local means denoising provides a slightly more sophisticated method where a pixel’s value is replaced by a mean which is sampled from other regions of the image. These sampled regions are only utilized if their mean is similar to the target region. This technique can preserve local texture information.

Single-cell segmentation SPEX includes several algorithms for single-cell segmentation. This includes traditional watershed cell segmentation in addition to a collection of pretrained deep learning models which include Stardist^25^ and Cellpose^18^. These algorithms will operate on the channel or combination of channels containing cell nucleus information. In instances where the input data differs in resolution to the training dataset for the respective models, the user can up-sample or downsample the image pixels. All segmentation modules will return an instance segmentation label image where each cell is assigned a unique integer value.

The optional post-processing modules aim to modify the single-cell segmentation labels. Tissue-based imaging is often prone to include technical artifacts. This could be tissue tears, folds or debris. These artifacts may generate false positive cell segmentation labels. Therefore, we include rule-based functions to exclude segmented labels based on morphological features.

In some rare instances, the pretrained deep learning segmentation may include false negative regions. This is likely a result of the model not having representation of this cell type in the training data. In these cases, one can utilize the optional rescue cells() algorithm . First, prototypical cell size and intensity are calculated adaptively from the currently segmented cell objects. These parameters are then used as input for classical watershed segmentation of the image. If a label from the watershed segmentation does not overlap with that of the deep learning approach, then it is merged into the final instance segmentation label image.

The user then has an option to expand the boundaries of the cell. This may be desired since some of the cell segmentation modules utilize only nuclear information. To ensure cytoplasmic compartments are included in feature extraction, boundaries should be dilated.

SPEX utilizes the scikit image regionprops function to extract single cell features. Currently, these features include mean intensity for each channel, extracted for each cell. The output will be a cell by expression matrix in Anndata formart which serves as input for downstream cell type clustering.

#### Proteomics Data Clustering

An Anndata object containing the cell by expression matrix obtained from the upstream image processing modules can be clustered into groups of cells that share the same properties. The SPEX proteomics clustering module leverages Phenograph for cell typing^14^. Within this module, users will define a transformation and scaling method that is most suitable for their data. Arcsin and Log available as transformation options. Winsorizing and z-scoring are available as scaling options.

#### Transcriptomics Data Clustering

Cells described by a cell by expression matrix obtained from the set of image processing or spatial transcriptomics ingestion modules can be clustered by gene expression. Currently, we provide a standard method to cluster the cells using the popular scanpy package. Cells with fewer than 20 total UMI counts were excluded from analysis. The cell by expression matrix is first normalized to the median total counts and log-transformed. The matrix is then whitened and PCA is performed (the optimal number of components is determined from the singular value spectrum). A nearest-neighbor graph 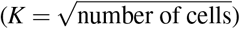 is constructed on the PCA matrix and the graph is clustered using the Leiden algorithm with the default parameters.

#### Spatial Analytics Module

##### Cell-cell co-occurrence at the cell population level

To systematically assess the non-random co-occurrence/avoidance of identified cell types at the cell population level, we used a permutation test to compare the number of interactions between all of the cell types in a given image (or in a given user defined region) to that of synthetic matched controls generated under the null hypothesis containing randomized cell phenotypes. The neighborhood radius (e.g. 20 µm) was chosen empirically to restrict cell-cell co-occurrence assessment to only those cells that are close enough to potentially establish physical contact between their membranes. This approach, similar to the one described in the histoCAT paper^26^, allows to determine the significance of cell-cell co-occurrence between two cell types.

##### Colocation Quotient: cell-cell co-occurrence at enhanced resolution

The Colocation Quotient (CLQ), defined as a ratio of ratios, is a widely used geographic index^27^. In the context of tissue images, it is used to measure the local density to the global density of a specific cell type^28^. Specifically, the local density is calculated as the proportion of cell type B in the neighborhood constructed by a certain radius, centered around cell type A. The global density is the proportion of cell type B in the entire tissue image. In general, if the density of cell type B within the neighborhood of A is more than the global density of B, the CLQ will be > 1. If the neighborhood of cell type A contains many other cell types other than B, the CLQ will be < 1. The value 1 means there is no spatial relationships between the two cell types.

The CLQ can be measured globally for a particular cell type A, CLQ_*A*→*B*_, or locally for each cell in the slide classified as that particular cell type, which we denote as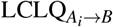. The CLQ is calculated through the following expressions:

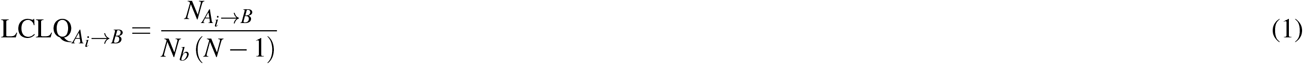

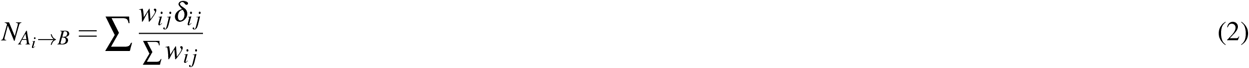

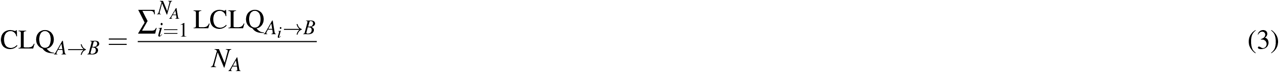

Where *N* is the total number of cells in the image, *δ*_*ij*_ is the Kroenecker delta indicating whether cell *j* is a type B cell; *w*_*ij*_ is 1*/N* for non-weighted version, and Gaussian distance decay kernel for weighted version which is common in geostatistics application li2011worldwide. The global CLQ will then be the average of the local CLQ.

##### Spatially informed transcriptomics using CLQ scores

For each cell type we estimated a CLQ score that describes the spatial relationship between cells (or globally as clusters), classifying them as “avoidant” or “attractive” spatial relationships. The cell-level CLQ classifications can be further investigated to identify differences in gene expression associated with for example “avoidant” behavior in clusters of interest. Gene expression differences were estimated using the scanpy (version 1.9.4)^29^ using default parameters. Hypothesis-driven testing of cell clusters and cell-cell spatial relationships for differences in gene expression can be performed flexibly using SPEX.

##### Spatially informed transcriptomics using SPEX clusters

For each cell in the dataset, neighborhood compositions were determined by collecting the counts of cells belonging to the differ-ent clusters included within a circle of radius epsilon centered on each cell (a parameter that can be adjusted by the user to assess the locality of expected cell interactions). SPEX communities then were identified via clustering of the matrix of the neighborhood compositions using Phenograph.

MERFISH single cell data was processed using the squidpy package^10^ (version 1.3.0) and then fed into SPEX for spatial analyses. For each cluster identified, CLQ was calculated and neighborhood compositions were determined for each cell. SPEX communities were identified via clustering neighborhood compositions using the Leiden algorithm as implemented in scanpy (version 1.9.4). These communities were then analyzed using decoupleR^30^ (version 1.4.0) to infer cell-cell interactions and pathways specific for said communities. SPEX communities were also inspected for pathway activities from a collection of well understood transcriptional pathways using progeny^31^ (version 1.12.0).

### SPEX Backend

SPEX’s architecture design is mainly driven by the requirements to the system performance, availability, and scalability. In particular, we aimed for an architecture supporting multi-user access to computing resources while ensuring and guaranteeing the execution of the resource-heavy tasks set by the user in a reasonable time frame. As an additional requirement, we considered the ease of system extension and maintenance. All these requirements are fulfilled by the microservice architecture that is inherently flexible as it is based on the idea of interoperability of small, loosely coupled and easily modifiable modules (microservices). A microservice is a small program that performs a clearly assigned task. Each microservice is hosted in a Docker container and can run individually or as a group via docker-compose. SPEX microservices communicate through the data bus instrumented by Redis). Redis manages message transfer between the ser-vices ensuring messages order and guaranteed delivery even if temporary system outages occur. Message distribution and load balancing mechanisms allow to easily increase the number of the microservices’ instances and spread them across different servers enabling straightforward scaling and performance boosting possibilities.

The integration of SPEX and the OMERO image management applications is implemented via a separate microservice. This service is responsible for creating OMERO sessions operating on two different layers.

The first layer, known as Python Blitz Gateway, is primarily used for direct access to images. It enables batch downloading of images in chunks (through ms-image-loader) and also permits the retrieval of image metadata not available through another layer of OMERO.The second layer, named OMERO WEB API, facilitates user interaction with images and projects. More specifically, it enables the request of metadata pertaining to user projects, accessible images, and compressed image fragments.Both layers are integral to SPEX since neither possesses all the necessary OMERO functions. Thus, to accomplish specific tasks, both layers must be used. They are created and managed by the microservice named ms-omero-session, and subsequently stored as objects in REDIS. From there, they can be accessed by other microservices and the backend system.

A proxy connection has been implemented to enable interactions with the OMERO WEB API. This mechanism proxies all requests to the OMERO WEB through our backend, utilizing a web session created on the OMERO WEB API layer by the ms-omero-session microservice. That session is then stored as an object in REDIS. This approach allows for requests to be overridden and reused, with the ms-omero-session microservice ensuring that the session remains active and accessible when necessary.

During the initiation of image processing scripts from the pipeline, the primary layer invoked is the Python Blitz Gateway. This layer is instantiated by the microservice referred to as ms-omero-session and is then stored as an object within REDIS.The ms-job-manager microservice necessitates a Python Blitz Gateway session. This session, created and stored in REDIS by the ms-omero-session microservice at the time of user login, is persistently monitored and refreshed to ensure its availability for reuse when required.

During the execution of pipeline blocks within the microservice ms-job-manager, the computational results are converted into the anndata-zarr format, which is compatible with Vitessce (https://github.com/vitessce/vitessce/). These converted data are then stored in a network storage system. This storage serves as a data source for subsequent use on the client’s frontend. The combination of Vitessce with data collection for creating interconnected datasets (multiple zarr files) into a unified dashboard enriches progressively with data from each pipeline step. This provides an excellent opportunity for a comprehensive overview of the entire research subject without the need to remember the full context of the experiment.

In SPEX, the data visualization system Vitessce is utilized, shifting data visualization workload from server to client side. This approach leverages cloud access capabilities within Vitessce. A key feature is the use of the h5ad to zarr format conversion, which is compatible with Vitessce. This conversion allows for efficient data handling, as it enables users to load only the data chunks currently in use, rather than the entire dataset. This not only reduces the load on the user’s device but also allows for repeated reuse of data obtained from pipeline outputs, with added filters and detail levels.

Furthermore, Vitessce is built upon WebGL2, which facilitates the transfer of data visualization processing to the graphics processing unit (GPU). Currently, Vitessce configuration files are backend-generated, offering simpler data visualization compared to seaborn and plotly. This requires initial technical expertise but ultimately reduces time in system support and development.

## Author contributions statement

X.L., X.P., C.F. T.R., D.O., and R.J. wrote the manuscript with input from all authors. X.L., X.P. , D.O. and R.J. conceived the SPEX platform. D.O. and R.J. supervised the project (D.O. from 2020 to 2022, and R.J. and J.G. 2022-present upon D.O.’s departure from Genentech). J.G. also provided software deployment strategy. R.J. designed the UI with input from authors and end users, developed image processing methods and executed image processing on spatial proteomics datasets. X.L., X.P., C.F., D.O. developed spatial analysis methods. X.P., D.H., and C.F. developed code for spatial transcriptomics analysis and executed analysis of spatial transcriptomics datasets. C.C. developed and tested single-cell data structure schema. T.R. analyzed and interpreted MIBI data. S.R., P.C., J.Z. executed method development of IMC PDAC data. D.O. performed clustering analysis of IMC Tonsil data. F.P. analyzed and interpreted IMC Tonsil spatial data. Z.S , M.N., and X.Y. generated Lung MERFISH data and executed primary analysis. L.M. provided interpretation and validation of spatial methods. J.C. and J.S. provided infrastructure engineering and OMERO integration work. H.C.B. provided software development supervision. A.Z. and E.T. executed frontend and backend software development. A.P. executed frontend and backend software development in addition to software testing. Both X.L. and X.P. contributed equally and have the right to list their names first in their CVs. All authors read and approved the final manuscript.

## Availability of data and materials

The open-source SPEX analysis platform is made publicly available at https://github.com/Genentech/spex Datasets used and/or analyzed in this work are made available on Zenodo

## Competing interests

X.L., X.P., D.O. and R.J. are co-inventors on a provisional patent application filed by Genentech/Roche relating to this manuscript.

## Author Affiliations

All authors were employees of Genentech at the time of project contributions. D.O.’s current affiliation is Cell Signaling Technologies. X.P.’s current affiliation is Revolution Medicines.

A.Z. and E.T. were contract employees at time of project contribution. A.Z.’s current email address is and E.T.’s current email is. All other authors are current employees of Genentech.

## Supporting Information

**Table 1.** Antibodies, conjugates, clones and working concentrations used in tonsil multiplex staining.

| Antibody | Conjugate | Clone | Concentration ($\mu\text{g/mL}$) |
| --- | --- | --- | --- |
| CD20 | 161-Dy | H1 | 3 |
| CD45 | 152-Sm | D9M8I | 7 |
| CD68 | 159-Tb | KP1 | 0.1 |
| CD8 | 162-Dy | CD8/144B | 2 |
| Collagen type I | 169-Tm | Polyclonal | 1 |
| Cytokeratin | 176-Yb | AE1/AE3 | 0.8 |
| Histone H3 | 171-Yb | D1H2 | 0.5 |
| Ki67 | 168-Er | B56 | 3 |
| BCL2 | 146-Nd | EPR17509 | 7 |
| CD25 | 175-Lu | EPR6452 | 5 |
| CD31 | 151-Eu | EPR3094 | 5 |
| CD336/Tim3 | 154-Sm | D5D5R | 7 |
| PD1 | 165-Ho | EPR4877 | 5 |
| PDL1 | 150-Nd | 73-10 | 10 |

## Notes

### Summary of Updates

Manuscript content changes and author list updated

